# Rapid spread of a vertically transmitted symbiont induces drastic shifts in butterfly sex ratio

**DOI:** 10.1101/2024.02.24.581844

**Authors:** Mai Miyata, Masashi Nomura, Daisuke Kageyama

**Affiliations:** Faculty of Engineering, University of Fukui, Fukui 910-8507, Japan; Life Science Innovation Center, University of Fukui, Fukui 910-8507, Japan; Graduate School of Horticulture, Chiba University, Matsudo, Chiba 271-8510, Japan; Institute of Agrobiological Sciences, National Agriculture and Food Research Organization, Owashi, Tsukuba, Ibaraki 305-0851, Japan

**Author notes:** Correspondence (M.M.), (M.N.), (D.K.).

## Abstract

Sex ratio dynamics constitutes a pivotal subject in evolutionary biology^1^. Under conditions of evolutionary equilibrium, the male-to-female ratio tends to be approximately 1:1; however, this equilibrium is susceptible to distortion by selfish genetic elements exemplified by driving sex chromosomes and cytoplasmic elements^2,3^. While previous studies have substantiated instances of these genetic elements distorting the sex ratio, studies specifically tracking the process with which these distorters spread within populations, leading to a transition from balanced parity to a skewed, female-biased state, are notably lacking. Herein, we present compelling substantiation regarding the rapid spread of the cytoplasmic endosymbiont *Wolbachia* within a localized population of the pierid butterfly *Eurema hecabe* (Figure 1A). This resulted in a shift in the sex ratio from near parity to an exceedingly skewed state overwhelmingly biased toward females, reaching 94.4% within a remarkably brief period of 4 years.

---

In *E. hecabe*, females fall into two categories (Figure 1B): a minority of females that possess a sex-ratio-distorting *Wolbachia w*Fem (CF females; Z0) and most females that lack this *w*Fem (C females; WZ)^4^. C females are infected with a non-sex ratio-distorting *Wolbachia w*CI and produce an equal ratio of male and female offspring after mating with *w*CI-infected males (C males; ZZ). In contrast, CF females are infected with a sex-ratio-distorting *Wolbachia w*Fem alongside *w*CI, resulting in female-only offspring after mating with C males^4^. In the sister species *Eurema mandarina*, a similar phenomenon is observed, where the prevention of maternal inheritance of the Z chromosome, coupled with the feminization of the resulting Z0 offspring, serves as the underlying mechanism for the exclusive production of females within CF lineages^5^.

**Figure 1.**
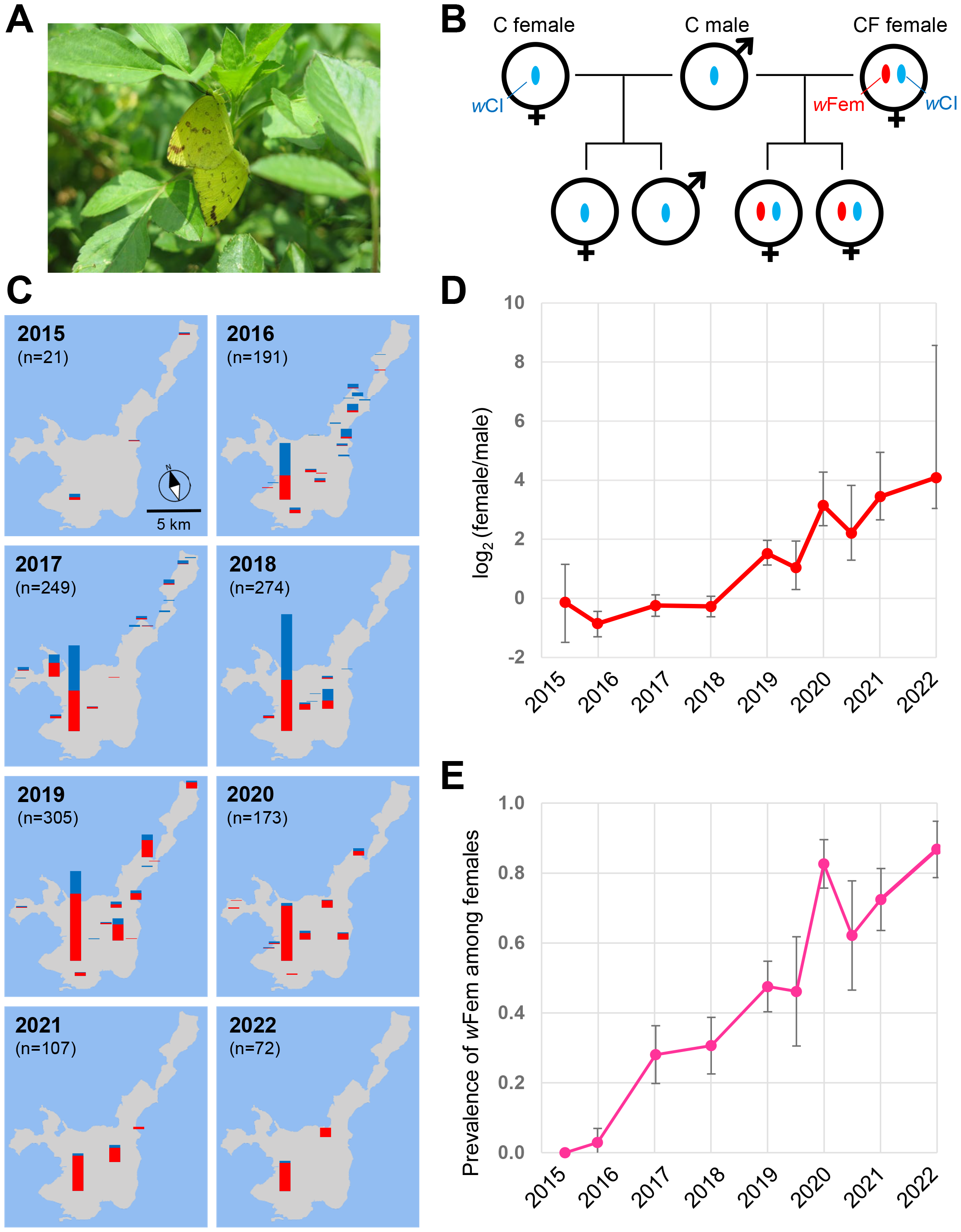
Sex ratios and *Wolbachia* infection frequencies of *Eurema hecabe* collected on Ishigaki Island between 2015 and 2022. (A) Field observation of a mating pair of *E. hecabe*. (B) Schematic illustration of the two maternal lineages (C and CF) of *E. hecabe*, with blue ovals indicating *w*CI and red ovals representing *w*Fem. (C) Temporal changes in sex ratios of *E. hecabe* collected from 32 locations on Ishigaki Island. (D) Female-to-male ratios (log_2_ female/male) with 95% confidence intervals. (E) Infection frequencies of *w*Fem (proportions of CF females among females) with corresponding 95% confidence intervals.

Based on a 2011 investigation, the frequency of CF females among the females in an *E. hecabe* population on Ishigaki Island was recorded at a relatively low frequency of 8.0% (4 CF females and 46 C females)^4^. Subsequently, our comprehensive collection effort yielded a total of 1,392 individuals of *E. hecabe* comprising 856 females and 536 males. This sampling was conducted across 32 sites on Ishigaki Island during nine visits from 2015 to 2022. The observed female-to-male ratio expressed as log_2_ (female/male) exhibited variability within the range of −0.86 (35.6% female) and −0.14 (47.6% female) 2015–2018. However, after 2019, a discernible increase in females occurred, elevating the ratio to 4.09 (94.4% female), as depicted in Figures 1C and 1D. Furthermore, the infection frequency of *w*Fem increased from 0.00–0.08 in 2015/2016 to 0.87 in 2022 (Figure 1E). Interestingly, the frequency of CF females and the female-to-male ratio tended to be lower during autumn; this was potentially attributed to heightened summer heat stress in the subtropical climate zone of Ishigaki Island. Under laboratory conditions, the offspring of 10 female butterflies gathered in April 2019 were reared to adulthood. Consistent with the findings of a previous study^4^, the offspring of seven females (CF females) exhibited an exclusively or predominantly female composition (157 females and 1 male; Supplementary Data). Conversely, the offspring of three C females comprised both males and females (87 females and 68 males; Supplementary Data). Thus, the rapid increase in the frequency of CF females appears to explain the prompt alteration observed in the sex ratio on Ishigaki Island.

The alteration in the population sex ratio has also been documented in the nymphalid butterfly *Acrea encedon*. The female ratio experienced a transition from 36.0% in 1909–1912 to an overwhelming 98.4% in 1963–1964 (estimated to have occurred within a span of 100–150 generations^6^). In subsequent research, the presence of male-killing *Wolbachia* was reported in this species^7^. In contrast, in *E. hecabe*, the substantial sex ratio shift potentially transpired more swiftly within an estimated 40 generations during 2018–2022. This projection is based on the assumption that *E. hecabe* experience ten generations per year on Ishigaki Island (Supplementary Data). The non-linear regression analysis based on the observed spread rate of *w*Fem suggested that the relative fitness of CF females is 0.54 (Supplementary Data). This means that CF females produce 1.07 times the number of C females. The rapid spread rate of *w*Fem within the *E. hecabe* population is comparable to that observed in cytoplasmic-incompatibility-inducing *Wolbachia*^8^. In contrast to cytoplasmic incompatibility, the pronounced strong sex ratio bias results in a scenario where males exhibit a heightened selective advantage compared with females. In the nymphalid butterfly *Hypolimnas bolina* and the green lacewing *Mallada desjardinsi*, the nuclear-encoded suppressor of endosymbiont-induced male-killing demonstrated rapid spread, causing the strongly female-biased sex ratio to revert to a balanced 1:1 within five years^9^. Significantly, *E. hecabe* males positive for both *w*CI and *w*Fem (CF males) were identified in *E. hecabe* samples collected between 2019 and 2021 (n = 7), a phenomenon absent in older samples. Should the appearance of CF males serve as an indicator of the presence of a host nuclear suppressor of feminization, it suggests the potential for a swift restoration of the sex ratio of *E. hecabe* on Ishigaki Island in the coming years. Conversely, in the sister species *E. mandarina*, a high infection frequency of *w*Fem (ca. 80% or more) persists within an island population (Tanegashima Island) for a minimum of 12 years, consistently leading to a distortion in the sex ratio toward females^10^. Our study highlights that not only suppressors but also cytoplasmic sex ratio distorters can spread rapidly. This indicates the possibility of quick and repetitive cycles of an evolutionary arms race between the cytoplasm and nucleus, which may lead to a rapid evolution of sex-determining systems.

