## Supplementary material for "Rapid spread of a vertically transmitted symbiont induces drastic shifts in butterfly sex ratio": Document S1

**Supplemental Table S1. Sex ratio, egg hatchability, and adult emergence rate of offspring derived from Ishigaki Island population in 2019.**

| Wild-caught mothers | *Wolbachia* infection status[1] | No. of eggs | No. of hatched larvae | No. of adults | | Egg hatch rate | Adult emergence rate[2] | Female ratio[3] |
|---|---|---|---|---|---|---|---|---|
| | | | | Female | Male | | | |
| 2019-231 | C | 166 | 92 | 36 | 30 | 0.55 | 0.40 | 0.55 |
| 2019-244 | C | 116 | 83 | 38 | 27 | 0.72 | 0.56 | 0.58 |
| 2019-248 | C | 61 | 38 | 13 | 11 | 0.62 | 0.39 | 0.54 |
| 2019-234 | CF | 74 | 22 | 13 | 0 | 0.30 | 0.18 | 1.00 |
| 2019-235 | CF | 24 | 6 | 5 | 0 | 0.25 | 0.21 | 1.00 |
| 2019-236 | CF | 33 | 27 | 12 | 0 | 0.82 | 0.36 | 1.00 |
| 2019-242 | CF | 85 | 51 | 28 | 0 | 0.60 | 0.33 | 1.00 |
| 2019-243 | CF | 83 | 35 | 22 | 0 | 0.42 | 0.27 | 1.00 |
| 2019-249 | CF | 187 | 65 | 54 | 0 | 0.35 | 0.29 | 1.00 |
| 2019-250 | CF | 101 | 35 | 23 | 1 | 0.35 | 0.24 | 0.96 |

[1]C: singly infected with *w*CI. CF: doubly infected with *w*CI and *w*Fem

[2]No. of emerged adults per no. of laid eggs.

[3]Proportion of females among adults.

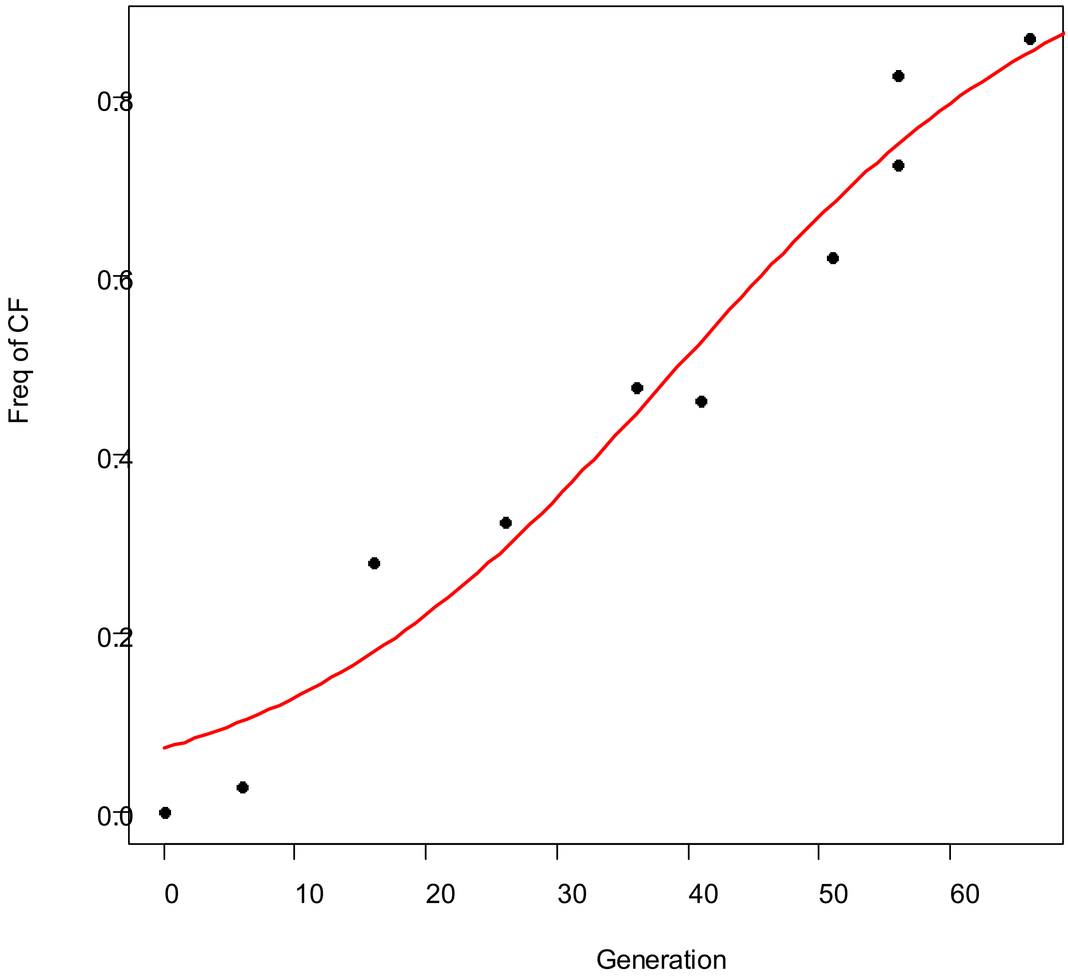


**Supplemental Figure S1. Non-linear curve fitting of field survey data.** Black plot: observed data. Red line: the fitted curve based on the model shown in the Supplemental Experimental Procedures.

### Supplemental Experimental Procedures

*Field sampling and laboratory rearing*

In 2015-2022, adults of *E. hecabe* were collected from 32 sites on Ishigaki Island, Japan (Fig. 1 A). Ten female butterflies collected in April 2019 were individually allowed to oviposit on the host plant *Ormocarpum cochinchinense* (Lour.) Merr. (Fabaceae). The collected eggs were separately reared in plastic containers until adult emergence under 16:8 h light-dark cycles at 25 °C. The hatched larvae were fed on an artificial diet containing leaf powder of *Albizia julibrissin* Durazz. (Fabaceae) described by Kato and Sakakura (1994)[1]. The sex ratio of emerged adults was recorded for each brood.

*DNA extraction and PCR*

Adult legs homogenized in 100 µl of STE buffer (100 mM NaCl; 10 mM Tris-Cl, pH 8.0; 1 mM EDTA) containing 2 µl of proteinase K (20 mg/ml) were incubated at 37 °C for 30 min and at 95 °C for 5 min. After centrifugation at 20,000 g, the supernatant was used as a template of PCR reactions. Infection with *Wolbachia* of either *w*CI or *w*Fem strain was identified by specific PCR detection targeting the *wsp* gene[2]. Specifically, *w*CI was detected by using the *Wolbachia*-specific forward primer wsp81F (5'–TGGTCCAATAAGTGATGAAGAAAC–3')[3] and *w*CI-specific reverse primer WHecFem1 (5'–ACTAACGTTTTTGTTTAG–3')[2], which amplify a 232-bp DNA fragment and *w*Fem was detected by using the *w*Fem-specific forward primer WHecFem2 (5'–TTACTCACAATTGGCTAAAGAT–3')[2] and *Wolbachia*-specific reverse primer wsp691R (5'–AAAAATTAAACGCTACTCCA–3')[3], which amplify a 398-bp DNA fragment. The temperature profile adopted for both PCR reactions are 35 cycles of denaturation at 95 °C for 1 min, annealing at 58 °C for 1.5 min, and extension at 72 °C for 1.5 min, followed by a final extension at 72 °C for 7 min. To discriminate three *Eurema* species (*E. hecab*e, *E. mandarina* and *E. blanda*) inhabiting on Ishigaki Island, the extracted DNA was subjected to the diagnostic PCR targeting the *Tpi* gene[4]. *E. hecabe* was identified using the *E. hecabe*-

specific forward primer Eh6-F (5'–TGTGGCCTTCTGCCCTATTAAA–3') and the reverse primer Eh6-R (5'–ACAGGCAATGACCTTGAGTC–3'), which amplify a 375-bp DNA fragment. The PCR condition was 94.0 °C for 5 min, followed by 35 cycles of 94.0 °C for 30 s, 48 °C for 30 s, 72.0 °C for 30 s, and finally 72.0 °C for 7 min.

*Statistical analysis*

Under the assumption that (1) C females produce equal numbers of males and females and (2) traits other than fertility does not differ between C females and CF females, the proportion of females ($P_t$) at generation t can be described as

$$P_t = P_0 (2k)^t / \{P_0 (2k)^t + (1 - P_0)\}$$

where $P_0$ depicts the initial proportion of CF females among females and k depicts the relative number of offspring (fertility) produced by CF females compared to the number of offspring produced by C females. We used this model for the non-linear regression analysis to estimate the value of k by inputting the empirical values of $P_t$ observed in ten different time points (field surveys). The regression analysis was performed using R version 3.4.4 (R Core Team 2017). The codes are shown as follows.

```
gen <- c(0,6,16,26,36,41,46,51,56,66) #generations inferred from
sampling date
i_freq <- c(0,0.029411765, 0.280701754, 0.325966851, 0.475409836,
0.461538462, 0.826086957, 0.621621622, 0.724489796, 0.867647059)
#frequencies of CF females among females
test_dat <- data.frame(gen, i_freq)
fit <- nls(i_freq ~ 1 / (1 + ((1/p) -1) * ((2*k)^(-gen))), test_dat,
start=list(p=0.1, k=1.1)) #non-linear least square (regression
analysis)
summary(fit)
p <- coef(fit)["p"]; k <- coef(fit)["k"]
cat("estimated p:", p, "\n"); cat("estimated k:", k, "\n")
plot(test_dat$gen, test_dat$i_freq, pch = 16, xlab = "Generation",
ylab = "Freq of CF",las=1)
a <- function(gen) {1 / (1 + ((1/p) -1) * ((2*k)^(-gen)))}
curve(a, 0,80, col = "red", lwd = 2, add = TRUE)
```

```
legend("bottomright", legend = "Fitting curve", col = "red", lty = 1,
cex = 0.8)
```

*Estimation of voltinism of E. hecabe on Ishigaki Island*

Based on the published data on the development from egg hatching to adult emergence[5], we estimated the developmental zero as 9.62 °C and the effective cumulative temperature during larval and pupal stages as 415.38 degree-days (°C d). The eggs usually hatch 3 days after oviposition at 25 °C (M. Miyata, personal observation). Besides, the adult females produce mature eggs 5 days after emergence at 25 °C[6]. Considering these factors, we made a rough estimate for the effective cumulative temperature from the egg to the next generation as 538.42 °C d (i.e., 415.38 + (25 - 9.62) $\times$ 8). Averages of yearly temperature for 1991 to 2020 on Ishigaki Island were obtained from the Japan Meteorological Agency [http://www.jma.go.jp/jma/indexe.html]. The voltinism was calculated as 10.0873 generations per year by dividing the cumulative degree-days by the effective cumulative temperature from the egg to the next generation (538.42 °C d).

**Acknowledgements**

We thank Takuya Yamaji and Hiroshi Arai for collecting butterflies. This work was supported by a JSPS KAKENHI grant (18J21090), Sumitomo Foundation for basic Sciences and Mitsubishi Foundation Grant in the Natural Sciences for young researchers to M.M. and a JSPS KAKENHI grant (23H02229) to D.K.

**Author contributions**

M.M., M.N. and D.K. designed the study. M.M., M.N. collected and analyzed the data. M.M., M.N. and D.K. wrote the manuscript. All authors read and approved the final manuscript.

**Declaration of interests**

The authors declare no competing interests.
